# Mechanical feedback enables catch bonds to selectively stabilize scanning microvilli at T-cell surfaces

**DOI:** 10.1101/527903

**Authors:** Robert H. Pullen, Steven M. Abel

## Abstract

T cells use microvilli to search the surfaces of antigen-presenting cells for antigenic ligands. The active motion of scanning microvilli provides a force-generating mechanism that is intriguing in light of single-molecule experiments showing that applied forces on stimulatory receptor-ligand bonds increase their lifetimes (catch-bond behavior). In this work, we introduce a theoretical framework to explore the motion of a microvillus tip above an antigen-presenting surface when receptors on the tip stochastically bind to ligands on the surface and dissociate from them in a force-dependent manner. Forces on receptor-ligand bonds impact the motion of the microvillus, leading to feedback between binding and microvillar motion. We use computer simulations to show that the average microvillus velocity varies in a ligand-dependent manner, that catch bonds generate responses in which some microvilli almost completely stop while others move with a broad distribution of velocities, and that the frequency of stopping depends on the concentration of stimulatory ligands. Typically, a small number of catch bonds initially immobilize the microvillus, after which additional bonds accumulate and increase the cumulative receptor-engagement time. Our results demonstrate that catch bonds can selectively slow and stabilize scanning microvilli, suggesting a physical mechanism that may contribute to antigen discrimination by T cells.

## 1 Introduction

T cells directly engage antigen-presenting cells (APCs) to search for surface-displayed antigens. They use the T-cell receptor (TCR) complex to discriminate between self and foreign ligands in the form of peptides presented by major histocompatibility complex (pMHC) molecules on the APCs. T cells are able to recognize small numbers of antigens amongst a vast number of self-pMHCs (1–3) while being sensitive enough to distinguish between peptides with a single amino acid difference (4, 5). However, a comprehensive understanding of the mechanisms governing antigen recognition remains elusive, and growing evidence points to the importance of mechanical forces at the T cell-APC interface (6–8).

Recent experiments have revealed two intriguing physical mechanisms related to antigen recognition: (i) T cells use microvillar protrusions to actively search APCs, suggesting a mechanism to scan large portions of the APC surface (9). (ii) The average lifetime of a bond between a TCR and an antigenic pMHC is maximized when there is an applied force on the TCR-pMHC complex (10, 11). Taken together, these results suggest feedback between microvillar motion and TCR-pMHC binding: Forces imparted by microvillar motion influence TCR-pMHC lifetimes, while tensions on individual TCR-pMHC complexes impact microvillar motion. To our knowledge, this feedback and its consequences have not been explored before.

Microvilli are fingerlike membrane protrusions that have been observed on T cells in a variety of studies (7, 9, 12–15). They also contain large numbers of highly localized TCRs (13, 14), and a recent study using lattice light-sheet microscopy revealed that T cells use microvilli to actively scan the surfaces of APCs (9). Interestingly, this study showed that cognate pMHC on the APC resulted in some microvilli becoming “stabilized,” with long-lived contact at localized regions on the APC surface. This stabilization occurred even when early intracellular signaling through the TCR was disrupted, suggesting that a physical mechanism might be responsible. Taken together, these studies suggest that microvilli play a role in antigen discrimination during early stages of T-cell activation.

Physical contact between the T cell and APC leads to mechanical forces at the cell-cell interface. TCR-pMHC complexes experience forces that arise from a variety of sources including cell motion, membrane undulations, active cytoskeletal processes, and the microvillar motion described above (7, 16). A number of recent experimental studies have characterized the force-dependence of individual TCR-pMHC dissociation kinetics (6, 10, 11, 17–21). When bound to stimulatory pMHC, the TCR exhibits catch-bond behavior, in which the average lifetime is maximized when a force ∼10 to 20 pN is applied to the protein complex. In contrast, a TCR bound to a nonstimulatory pMHC exhibits slip-bond behavior, in which the lifetime strictly decreases with increasing force. The ligand- and force-dependence of the dissociation kinetics has been proposed as a potential mechanism to enhance discrimination between self and foreign pMHC (10, 22).

In this paper, we investigate the mechanical coupling between microvillar motion and TCR-pMHC binding. We are interested in whether the interplay between the two provides a physical mechanism that could impact antigen recognition. To this end, we introduce a physically-motivated theoretical framework describing the motion of a microvillus near an antigen-presenting surface. The framework captures key biophysical features while being simple enough to analyze in detail. When possible, we use experimentally-derived parameters, including those for force-dependent TCR-pMHC dissociation kinetics. In the results, we first characterize the motion of scanning microvilli in the presence of surfaces containing nonstimulatory (slip) pMHC, stimulatory (catch) pMHC, and mixtures of the two. We characterize the distribution of velocities for different cases and assess when an individual microvillus has stopped. We then characterize the total time of receptor engagement as a proxy for the degree of stimulation of TCRs at the microvillus tip. We conclude by discussing some assumptions of the model, the physical picture that emerges from our simulations, and potential implications for antigen recognition by T cells.

## 2 Methods

We consider a theoretical framework in which an isolated microvillus scans across an antigen-presenting surface. The velocity of the microvillus depends on the forces exerted on the microvillus by TCR-pMHC complexes (“bonds”). The number of bonds, bond lifetimes, and microvillus velocity are dependent upon the interplay between stochastic binding events, diffusive processes, and forces that result from the stretching and compression of bonds by the moving microvillus. We use a stochastic reaction-diffusion framework that accounts for the microvillus motion and the forces on TCR-pMHC bonds.

### 2.1 Computational framework

Figure 1 provides a schematic depiction of the model. We represent a patch of the antigen-presenting surface as a rectangular domain in which pMHC molecules diffuse. The tip of the T-cell microvillus is represented by a circular surface with a diameter of 100 nm (13, 15, 23, 24). We assume that it resides at a fixed distance above the antigen-presenting surface, which we take to be the length of a TCR-pMHC complex (*z*_bond_). TCRs diffuse about the microvillus tip and bind to pMHC molecules on the antigen-presenting surface. TCRs and pMHCs are represented by particles with a 5 nm radius (25), and particles on the same surface cannot overlap due to excluded volume. When TCR-pMHC bonds are stretched relative to their natural length, they impose a force on the microvillus. We describe the force with a linear spring model, *f* = *k*_bond_(*L* – *z*_bond_), where *k*_bond_ is the spring constant, *L* is the distance between the two bound particles, and the force is directed along the bond.

**Figure 1:**
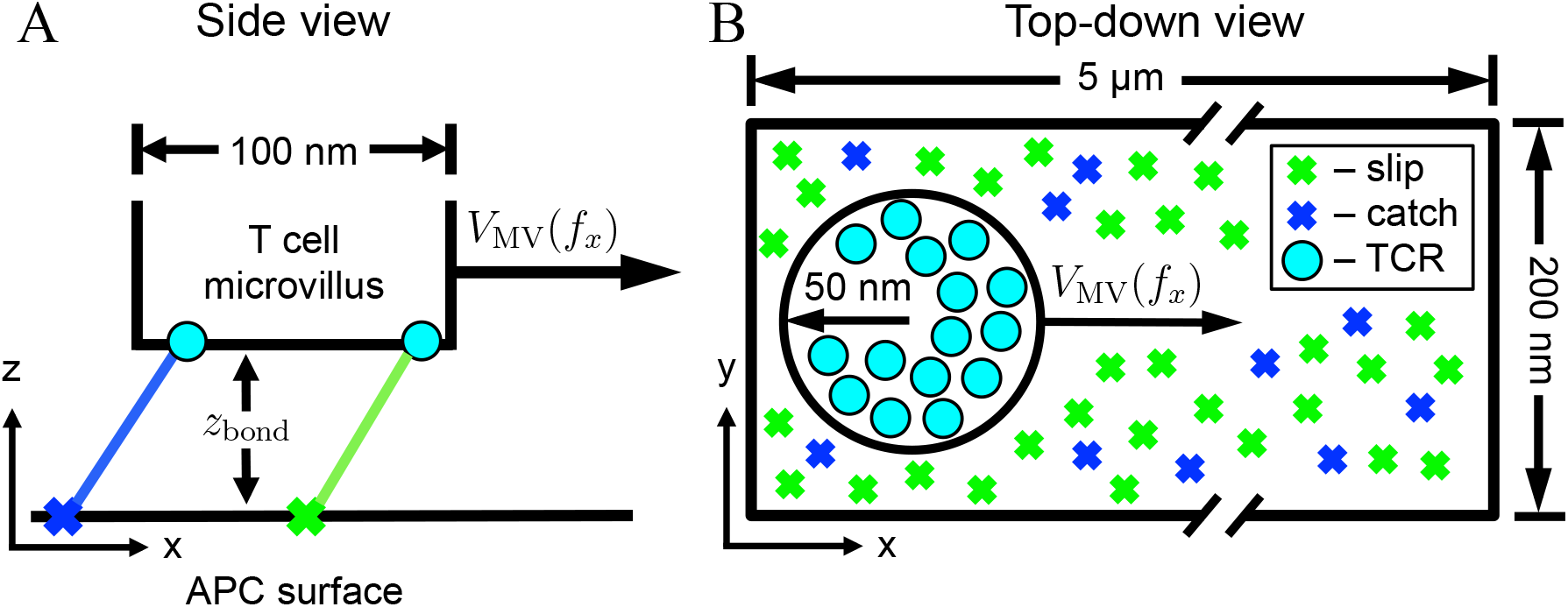
Schematic of a T-cell microvillus scanning across the surface of an antigen-presenting cell. TCR-pMHC bonds stochastically form and dissociate as the microvillus moves across the APC surface. The velocity, *V*_MV_, depends on the force exerted on the microvillus tip by TCR-pMHC complexes. (A) Side view of the system with the T-cell microvillus tip residing above the APC surface. (B) Top-down view. The microvillus moves in the *x*-direction. A mixed population of pMHC is shown; some form catch bonds upon binding TCRs while others form slip bonds.

The microvillus tip moves across the antigen-presenting surface in the *x*-direction. In the absence of an external force, the microvillus moves at velocity *V*_0_. However, forces arising from TCR-pMHC bonds impact the microvillus velocity, which we assume depends linearly on the component of the net force in the *x*-direction, *f_x_*(*t*):

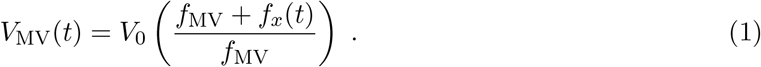

Here, *f_MV_* is a characteristic force and the time-dependence of *f_x_* is a consequence of the formation, stretching, and breaking of bonds. We assume that active processes driving the microvillus keep it in close apposition to the APC surface, and thus we neglect motion in the *z*-direction. Additionally, we ignore velocity fluctuations in the *y*-direction, as these would average to zero and can be thought of as changing the frame of motion of the microvillus. Physically, for an individual TCR-pMHC bond, when the pMHC is “in front of” the TCR (*x*_pMHC_ > *x*_TCR_), it effectively pulls the microvillus forward and increases the velocity. When the pMHC is “behind” the TCR, it decreases the velocity. The linear dependence of the velocity on force is consistent with the terminal velocity of an object experiencing a viscous drag. We discuss this choice and the results of a different functional form for *V*_MV_ later in the paper.

We use a discrete-time, continuous-space stochastic algorithm to describe the dynamics of the system. During each time step, the algorithm accounts for diffusive hops of particles, binding events, and dissociation of bonds. At the end of each time step, the position of the microvillus is updated in accordance with its velocity, which impacts the state of the system by changing the lengths of TCR-pMHC bonds and the positions of TCRs relative to the antigen-presenting surface. Details of the algorithm are presented in the SI.

Parameters used in the model are summarized in Table 1. The system size is 200 nm × 5 *μ*m, which is sufficiently long for the microvillus tip to remain within the domain. The total pMHC concentration is fixed at 100 pMHC/*μ*m^2^ (26–28), and the microvillus tip contains 23 TCRs (13). The state of the system is recorded every 2.5 × 10^−4^ seconds.

**Table 1:** Parameters used in the model.

| Variable | Definition | Value | Units | Source |
| --- | --- | --- | --- | --- |
| $D_{pMHC}$ | Diffusion coefficient, pMHC | $3.00 \times 10^{-3}$ | $[\mu\text{m}^2/\text{s}]$ | (29) |
| $D_{TCR}$ | Diffusion coefficient, TCR | 0.01 | $[\mu\text{m}^2/\text{s}]$ | (30) |
| $\Delta r$ | Diffusive step length | 10 | [nm] | — |
| $V_0$ | Microvillus velocity, no force | 5.20 | $[\mu\text{m}/\text{min}]$ | (13) |
| $f_{MV}$ | Threshold force, microvillus velocity | 50 | [pN] | — |
| $k_{\text{bond}}$ | Compressional stiffness of bonds | 1.00 | [pN/nm] | (16) |
| $k_{\text{nonstim}}^{\text{on}}$ | 2D on-rate, nonstimulatory pMHC | $2.6 \times 10^2$ | $[\text{nm}^2/\text{s}]$ | (17) |
| $k_{\text{stim}}^{\text{on}}$ | 2D on-rate, stimulatory pMHC | $2.4 \times 10^5$ | $[\text{nm}^2/\text{s}]$ | (17) |
| $z_{\text{bond}}$ | TCR-pMHC complex length | 13 | [nm] | (31) |
| $\Delta t$ | Time step | $2.5 \times 10^{-5}$ | [s] | — |

### 2.2 TCR-pMHC binding kinetics

In the model, TCR-pMHC dissociation kinetics are described by the Bell model for slip bonds (nonstimulatory pMHCs) and by the two-pathway model for catch bonds (stimulatory pMHCs). The off-rate in the Bell model is characterized by (32)

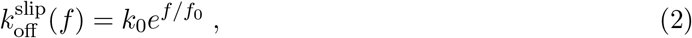

where *k*_0_ is the off-rate at zero applied force, *f* is the force on the receptor-ligand complex, and *f*_0_ is the reference force. The off-rate in the two-pathway model is characterized by (33)

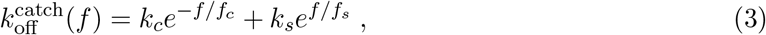

where *c* and *s* denote catch- and slip-phase parameters, respectively.

We parameterized experimental bond-lifetime data from two previous studies that showcase catch- and slip-bond binding characteristics. Liu et al. used a biomembrane force probe to characterize the interactions of the OT1 TCR with a panel of pMHCs that ranged from a nonstimulatory peptide (E1) to a strongly stimulatory peptide (OVA) (10). Feng et al. utilized opticals tweezers to investigate interactions of the N15 TCR with the stimularory vesicular stomatitis virus octapeptide (VSV8) presented by the MHC class I molecule H-2K^b^ (19). Figure 2 shows the average bond lifetime as a function of tensile force for the cases we consider in this paper. We also consider a hypothetical slip bond (“strong slip”) with the same maximum lifetime as a VSV8 catch bond and the same reference force as the E1 slip bond. The strong slip bond is used as a control to determine if a slip bond with a large zero-force lifetime behaves similarly to the catch-bond cases. For convenience, we refer to the two catch bonds and the strong-slip bond as “stimulatory.” Parameters for the different cases are tabulated in the SI (Table S1).

**Figure 2:**
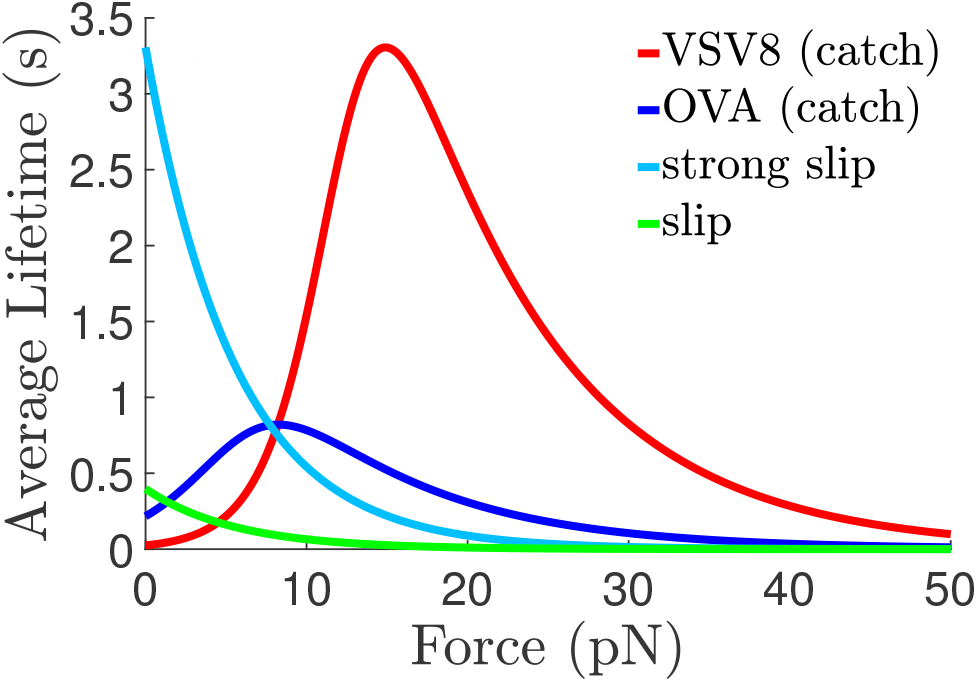
Average TCR-pMHC lifetimes as a function of tensile force. Curves for VSV8, OVA, and slip were fit using nonlinear least square fits of experimental data from Refs. (10) and (19). The “strong slip” bond is a hypothetical control with the same maximum average lifetime as VSV8 and the same reference force as the slip bond.

To characterize binding rates, we use data from Huang et al. (17), which reports effective 2D on-rates for the OT1 TCR binding to OVA and to E1 pMHC (Table 1). We use the on-rate reported for OVA to describe the binding of all stimulatory pMHC in this study (OVA, VSV8, and “strong slip”).

## 3 Results

In this section, we characterize the collective effects of catch and slip bonds on the motion of of microvilli. The total pMHC concentration is fixed at 100 *μ*m^-2^ throughout. For reference, we consider an antigen-presenting surface containing only nonstimulatory ligands with slip-bond kinetics. We then mimic varying degrees of stimulation by replacing 10%, 20%, 30%, and 100% of the nonstimulatory pMHCs with stimulatory pMHCs (either VSV8, OVA, or strong slip). For each condition, we generate 25 independent trajectories to assess stochastic effects.

### 3.1 Scanning microvilli are slowed in a ligand-dependent manner

Figure 3 shows the average microvillus displacement over time for the three different stimulatory pMHCs at various fractions of the total concentration. The green line (0% stimulatory pMHC) represents the average microvillus position when only nonstimulatory slip bonds are present. For this case, the average displacement grows linearly in time with a velocity of 3.5 *μ*m/min, which is less than the zero-force velocity of the microvillus (*V*_0_ = 5.2 *μ*m/min). From Eqn. 1, the average velocity is consistent with the nonstimulatory TCR-pMHC bonds exerting an average net force of approximately 16 pN in the direction opposed to the microvillus motion.

**Figure 3:**
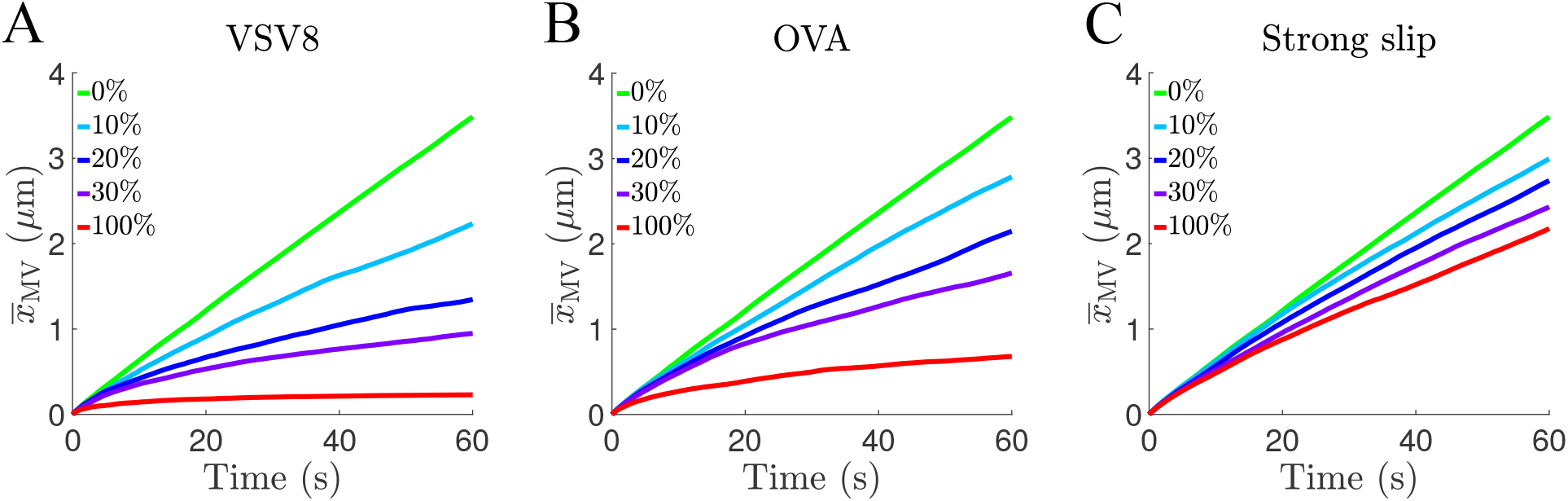
The average microvillus displacement for different fractions of stimulatory pMHC: (A) VSV8, (B) OVA, and (C) strong slip. Each line shows the average displacement from 25 independent trajectories.

The average displacement of the microvillus decreases as the fraction of stimulatory pMHCs increases. VSV8 leads to the most pronounced decrease in microvillus displacement, with increasing fractions of VSV8 leading to slower average microvillus movement. For 100% VSV8, the average displacement is nearly completely arrested after a short time. OVA generates similar behavior with smaller relative changes as the fraction of OVA is changed. The strong-slip case shows the weakest change in the average displacement as its fraction increases.

### 3.2 Heterogeneity of microvillus trajectories

Stochastic binding and dissociation events lead to differences in the motion of microvilli even at identical conditions. Figure 4 shows the time-dependent positions of microvilli from multiple independent simulations. When only nonstimulatory pMHCs are present (Figure 4A), there is relatively small variation between individual displacements and the average microvillus displacement. However, when the fraction of VSV8 pMHC is 10%, there is markedly more variation between individual microvillus displacements. One particularly striking feature is that some trajectories plateau for sustained periods of time. During these periods, the microvillus is approximately stationary due to forces exerted by TCR-pMHC bonds. When the fraction of VSV8 pMHC is 30%, a greater fraction of microvilli become effectively immobile within the one-minute period, which further decreases the average microvillus displacement. Analogous figures with OVA and strong-slip pMHC are shown in Figure S1.

**Figure 4:**
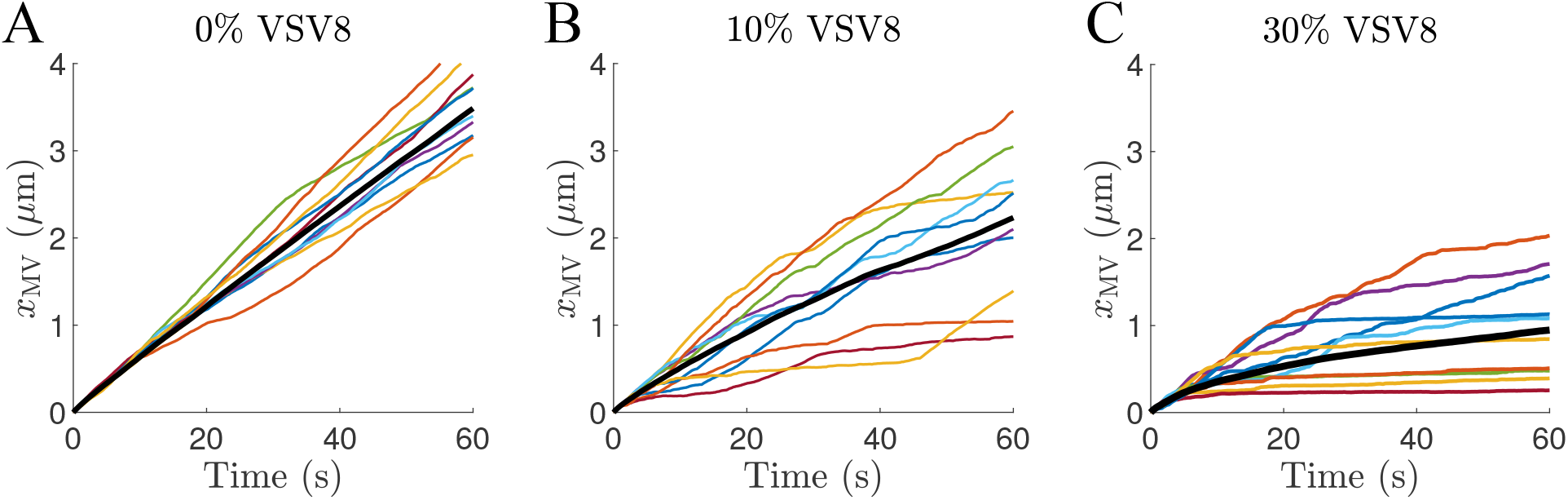
Displacements of microvilli for (A) nonstimulatory pMHC, (B) 10% VSV8 pMHC, and (C) 30% VSV8 pMHC. Black lines show the average microvillus displacement calculated from 25 independent trajectories. Colored lines show the displacement of individual microvilli (10 shown).

Movies S1 and S2 show sample trajectories at 10% and 30% VSV8 pMHC, respectively. In each, the microvillus experiences a period during which it is nearly stationary. For these cases, it can be seen that binding a small number of catch bonds significantly impacts the velocity of the microvillus. Once the catch bonds significantly slow the microvillus, additional slip bonds accumulate over time.

### 3.3 Catch bonds stabilize microvilli

Figure 5 characterizes distributions of the microvillus velocity for different types and fractions of stimulatory pMHC. When only nonstimulatory pMHCs are present (green line), the velocity distribution is approximately normal with a velocity of 3.48± 1.08 *μ*m/min (mean ± SD). Figure 5A shows how the presence of VSV8 pMHC changes the velocity distributions. At 10% VSV8, a small peak in the probability density emerges near 0 *μ*m/min, which is associated with stationary (“stabilized”) microvilli. As the fraction of VSV8 increases, both the number and duration of immobile microvilli increase, leading to more prominent and narrower peaks. Results for OVA (Figure S2) are similar but less pronounced. Figure 5B compares distributions of the velocity for systems containing varied fractions of strong-slip pMHC. Although larger fractions of strong-slip pMHC decrease the average velocity, in contrast with Figure 5A, the velocities extend across a broad range with no peak near 0 *μ*m/min. Thus, even large fractions of strong-slip pMHC do not result in the effective immobilization of microvilli.

**Figure 5:**
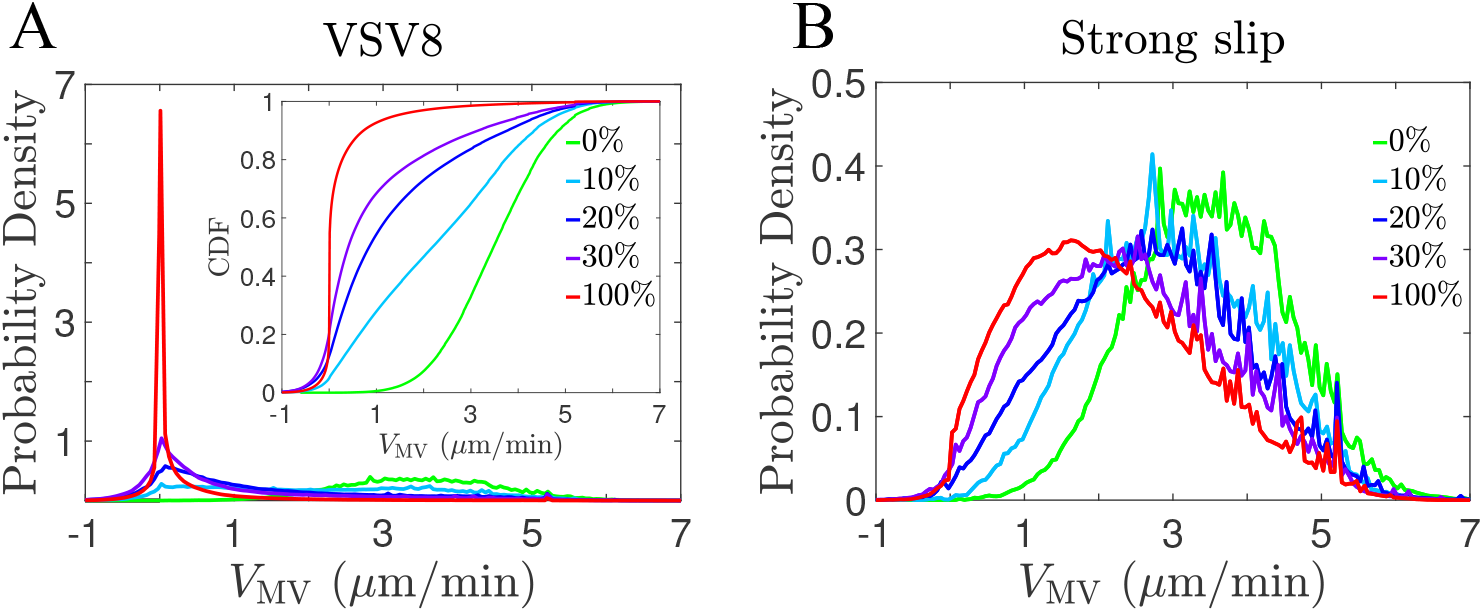
(A) Probability density and cumulative distribution functions (inset) of the microvillus velocity at various fractions of VSV8 pMHC. (B) Probability density of the microvillus velocity at various fractions of strong-slip pMHC. Each distribution is constructed from the velocities obtained over the course of 25 independent trajectories. The green curve (0%) is the same on both figures.

To further characterize the stabilization of microvilli by stimulatory pMHC, we determine the probability that a microvillus effectively stops during the course of a simulation (Figure 6). We define a “stopping event” to be when a microvillus has an average velocity ≤ 0.25 *μ*m/min for a continuous period of at least 10 s. Varying the thresholds for the average velocity and the time interval did not affect our conclusions. Figure 6 shows that the microvillus did not stop for any fraction of strong-slip pMHC. However, stopping does occur when stimulatory pMHC with catch bond characteristics (OVA and VSV8) are present. For OVA, stopped microvilli are observed at 20% OVA, and there is a significant increase in the number of stopped microvilli at larger fractions of OVA. For VSV8, stopped microvilli are observed at all fractions of VSV8, with larger fractions increasing the likelihood of stopping.

**Figure 6:**
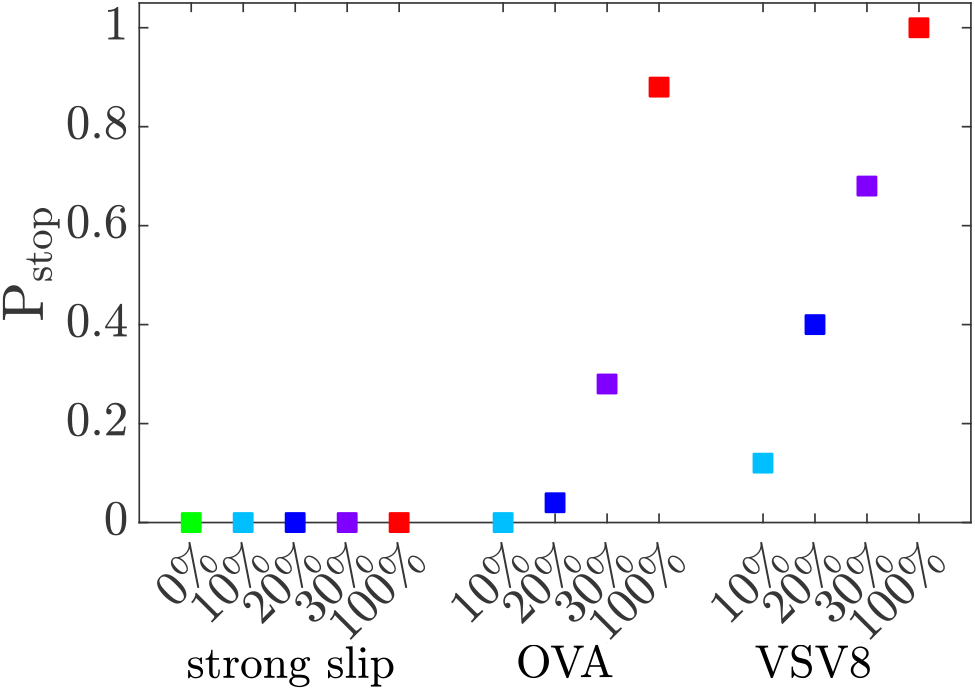
The probability that a microvillus is immobilized (“stops”) within 1 minute of scanning. A “stopping event” occurs when the average velocity is ≤ 0.25 *μ*m/min for a continuous period of at least 10 s. Each point is obtained from 25 independent trajectories.

### 3.4 Catch bonds impact cumulative TCR-pMHC binding times

In Figure 7, we examine the time-dependent displacement of a microvillus concurrently with the number of slip and catch bonds. Figure 7A shows a case with only nonstimulatory pMHC present. Although the number of slip bonds varies during the simulation, the slope of the microvillus displacement remains relatively constant. This is consistent with a rapid turnover of bonds, in which they frequently form and break. Figures 7B and 7C correspond to trajectories with 10% and 30% VSV8 pMHC, respectively. These two sample trajectories correspond to Movies S1 and S2. In both cases, when two or more catch bonds form, the microvillus slows dramatically. After the microvillus is initially stabilized, there is a gradual accumulation of additional slip bonds. In Figure 7B, the sudden increase in velocity at late times is coincident with the breaking of catch bonds and a rapid decrease in the number of slip bonds. Figure S3 shows the average number of catch and slip bonds as a function of the microvillus velocity. These results are consistent with TCR-pMHC bonds accumulating at stabilized microvilli: In systems with catch bonds, slower velocities have larger average numbers of TCR-pMHC bonds.

**Figure 7:**
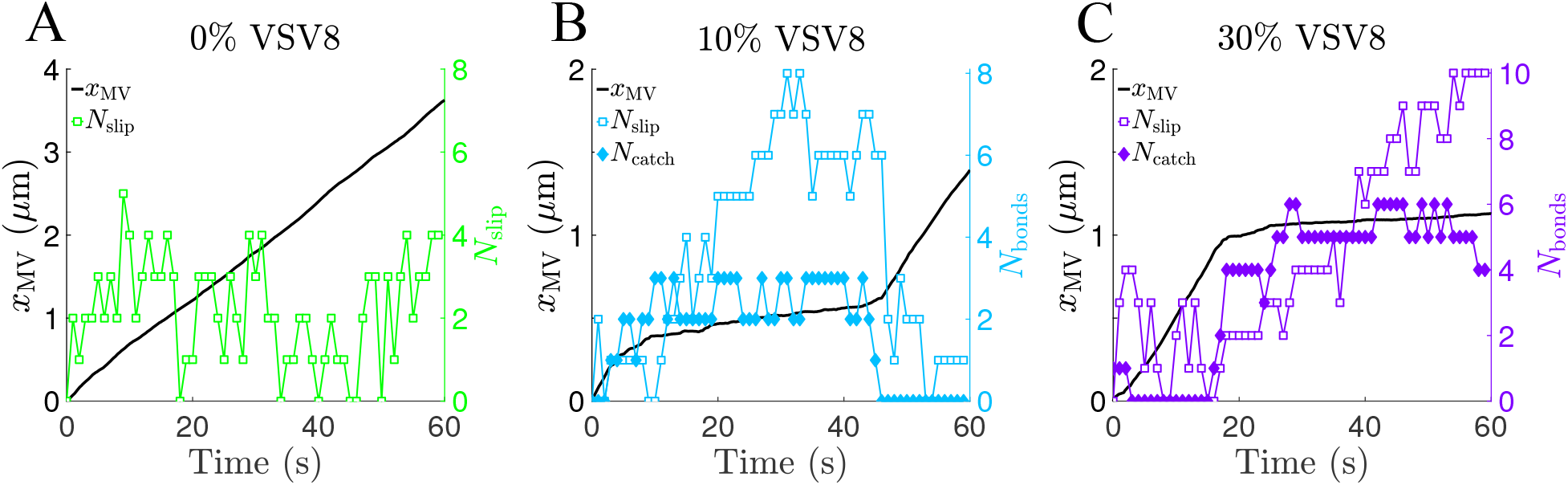
The number of catch and slip bonds (colored lines, right axes) and the microvillus displacement (black lines, left axes) for sample trajectories at three fractions of VSV8 pMHC (A-C).

Our results indicate that catch bonds selectively stabilize scanning microvilli in a stochastic manner and that TCR-pMHC bonds accumulate when the microvillus is stopped. This suggests a physical mechanism that could link the motion of the microvillus to intracellular processes through receptor engagement. As a proxy for the degree of stimulation through the TCR, we calculate the cumulative time of receptor engagement as a function of time for each trajectory:

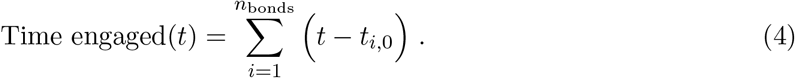

Here, *n*_bonds_ is the number of bound TCRs at time *t*, and *t*_*i*,0_ is the time of bond formation for a given TCR-pMHC complex. Thus, at time t, this function gives the total time that all current TCR-pMHC complexes have been present. Given the results on microvillus stopping in Figure 6, we analyze trajectories with and without stopping events separately to provide insight into the effects of microvillar stabilization by catch bonds.

When only nonstimulatory slip bonds are present, the average cumulative time engaged is 0.26 s. Figure 8A shows the average time of engagement for trajectories with a microvillus-stopping event for various fractions of VSV8 pMHC. The average time engaged increases with increasing fraction of stimulatory pMHC, exceeding 10 s for 30% VSV8. Note that these curves underestimate the typical time engaged for a specific stabilized microvillus because the average occurs over periods in which some microvilli are stopped and others are not. In comparison, Figure 8B shows a substantially smaller average time engaged for trajectories in which the microvillus does not stop. Figure 8C shows the average time engaged for systems with various levels of strong-slip pMHC, which did not induce any stopping events. The time engaged increases with an increasing fraction of strong-slip pMHC, but in all cases, the time engaged is substantially lower than corresponding cases in which catch bonds resulted in a stopping event (Figure 8A). Thus, systems with catch bonds are able to generate much longer sustained receptor engagement in comparison with even the strong-slip system.

**Figure 8:**
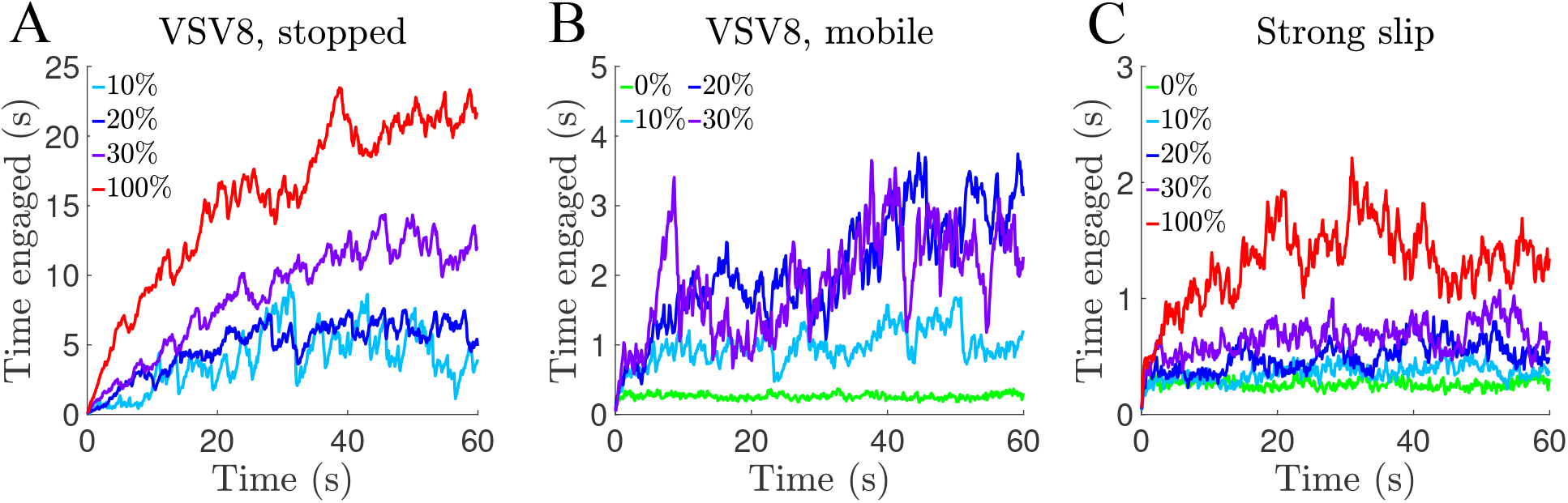
The average cumulative time of receptor engagement for various systems: (A) VSV8 with stopped microvilli, (B) VSV8 with mobile microvilli, and (C) strong-slip pMHC. No stopping events were observed for the strong-slip case.

## 4 Discussion

In this paper, we introduced a theoretical framework to explore the motion of a T-cell microvillus scanning across the surface of an antigen-presenting cell. TCRs at the tip of the microvillus interact with pMHCs on the antigen-presenting surface, and mechanical forces on the TCR-pMHC complexes resulting from motion of the microvillus impact both TCR-pMHC lifetimes and the microvillus velocity. Using parameters obtained from experimental studies of T cells, we showed that relatively small numbers of stimulatory pMHC with catch-bond characteristics immobilized microvilli in the model through mechanical coupling. In contrast, slip bonds alone did not immobilize microvilli. These results are exemplified by Figures 5 and 6, which show the distribution of velocities for scanning microvilli and the likelihood that a microvillus becomes immobilized. The effects become more pronounced as the concentration of antigenic pMHC increases.

### 4.1 Choice of the velocity profile, *V*_MV_

The mechanisms underlying microvillar motion on T cells have not been fully elucidated experimentally (9, 23, 34), so we chose to use a simple, physically motivated model relating the applied force to microvillar velocity. Our model uses a linear relationship between the velocity, *V*_MV_, and the applied force in the direction of motion. This relationship is consistent with the terminal velocity (zero acceleration) of an object experiencing a viscous drag: *F_A_* + *f_x_* – *μV*_M*V*_ = 0. Here, *F_A_* is a constant applied force driving the motion of the object (provided, for example, by cytoskeletal processes within the cell), *f_x_* is the applied force due to bonds, and *μ* is the drag coefficient relating the velocity to the drag force *μV*_M*V*_. Thus, *V_MV_* = (*F_A_* + *f_x_*)/*μ*, which is consistent with Eqn. 1.

The characteristic force in Eqn. 1, *f*_MV_, is unknown. If *f*_MV_ is too small, then even single slip bonds would significantly slow the microvillus. If *f*_MV_ is too large, then even a large number of bonds would not be able to significantly slow the microvillus. Both of these limits are inconsistent with the results from Ref. (9). With these limits in mind, we chose the value to be a few times larger than both typical forces experienced by individual TCR-pMHC bonds (35) and the force associated with the peak lifetime for catch bonds (10, 11). Moderately changing the value of *f*_MV_ impacts quantitative aspects of our results, although it does not impact the observation that catch bonds selectively stabilize microvilli. Figure S4 shows this for 2-fold changes in the force threshold for purely stimulatory and nonstimulatory pMHC systems.

Other models for *V*_MV_ may also be appropriate. Another plausible mechanism includes a “stall force” such that applied forces below a threshold have little effect on the velocity, while forces above the threshold have a large effect. To this end, we also consider a velocity profile with a Hill-like (sigmoidal) form in the SI (text and Figure S4). This profile results in an even more pronounced impact of catch bonds due to the inability of nonstimulatory ligands to significantly slow the microvillus tip on their own.

### 4.2 A physical mechanism for microvillar stabilization and enhanced antigen discrimination

T cells exhibit remarkable specificity and sensitivity in their search for antigenic ligands, yet the underlying mechanisms are still not fully understood. A growing body of work has revealed that physical mechanisms involving mechanical forces may play an important role (7, 8). In this paper, we have explored feedback between microvillar motion and TCR-pMHC dissociation kinetics. We were intrigued by the results from Cai et al. (9), which revealed that T cells use microvilli to scan the surfaces of APCs and that the presence of cognate pMHC stabilized microvilli even in the absence of tyrosine kinase signaling, which is a key component of the early TCR signaling pathway. In light of recent single-molecule studies of TCR-pMHC binding kinetics (10, 11), we hypothesized that nontrivial force-dependent dissociation kinetics could potentially lead to the stabilization. To investigate this idea, we developed a model that captures key biophysical properties of the microvillus-APC interaction but that is simple enough for analysis.

Taken together, our simulation results reveal a purely physical mechanism—the mechanical coupling of microvillar motion and TCR-pMHC binding—that could enhance antigen recognition by T cells. The physical picture is that forces generated by scanning microvilli provide a means to mechanically “test” TCR-pMHC bonds connecting the microvillus tip to the antigen-presenting surface. The total force necessary to immobilize a scanning microvillus is too large for an individual TCR-pMHC complex to reach before rupture. Thus, if only a small number of nonstimulatory TCR-pMHC bonds are present, the microvillus will continue to move, constantly increasing the force on the bonds and promoting their rupture. Short engagement times would prevent intracellular signaling through the TCR pathway.

In contrast, when a TCR binds to a stimulatory pMHC, the resulting catch bond is able to withstand substantially larger forces, thus allowing it to slow the microvillus and prolong the time spent in the range of forces that enhance its lifetime. For the VSV8 system, forces in the range of ∼10 to 30 pN significantly increase the lifetime of the bond. Thus, two such bonds aligned so that the force is mostly opposed to the direction of microvillus motion can halt the motion of the microvillus for a sustained period of time. This allows other TCR-pMHC complexes to accumulate and further stabilize the microvillus. As a consequence, the total time that TCRs stay engaged increases substantially when the microvillus is immobilized, which can promote intracellular signaling, starting with phosphorylation of ITAM domains on the cytoplasmic domain of the TCR complex (36, 37). Because the stimulatory pMHCs are randomly distributed, the time at which the microvillus stops occurs at random. Microvilli are immobilized more quickly on average at higher concentrations of stimulatory pMHC because of the increased likelihood to encounter multiple stimulatory pMHCs at once.

Our results reveal that the physical stabilization of microvilli could amplify differences in the response to self and foreign pMHCs, thus enhancing the ability of a T cell to identify antigens. Other potential advantages that microvilli confer to T cells are that they enable the scanning of large portions of the APC surface, facilitate contact between TCRs and pMHCs, and may provide protrusive and retractive forces (9, 38). Other molecules such as coreceptors, transmembrane phosphatases, and adhesion molecules are also important players in T-cell activation. This is a rich area for future study, and combining experiments and theory should help to untangle the complex mechano-chemical events leading to antigen discrimination by T cells. Additionally, it is interesting to speculate about active search and force generation as a tool in designing synthetic systems for detecting ligands of interest.

## 5 Acknowledgements

This work was supported by National Science Foundation CAREER Award PHY-1753017. It was conducted in part while RHP III was a Graduate Research Assistant at the National Institute for Mathematical and Biological Synthesis, an Institute sponsored by the National Science Foundation through NSF Award #DBI-1300426, with additional support from The University of Tennessee, Knoxville.

## References

1. Sykulev, Y., M. Joo, I. Vturina, T. J. Tsomides, and H. N. Eisen, 1996. Evidence that a single peptide-MHC complex on a target cell can elicit a cytolytic T cell response. Immunity 4:565–571.

2. Huang, J., M. Brameshuber, X. Zeng, J. Xie, Q. Li, Y. Chien, S. Valitutti, and M. M. Davis, 2013. A single peptide-major histocompatibility complex ligand triggers digital cytokine secretion in CD4+ T cells. Immunity 39:846–857.

3. Pageon, S. V., T. Tabarin, Y. Yamamoto, M. Yuangqing, P. R. Nicovich, J. S. Bridgeman, A. Cohnen, C. Benzing, Y. Gao, M. D. Crowther, K. Tungatt, G. Dolton, A. K. Sewell, D. A. Price, O. Acuto, R. G. Parton, J. J. Gooding, J. Rossy, J. Rossjohn, and K. Gaus, 2016. Functional role of T-cell receptor nanoclusters in signal initiation and antigen discrimination. Proc. Natl. Acad. Sci. U.S.A. 113:E5454–E5463.

4. Hogquist, K. A., S. C. Jameson, W. R. Heath, J. L. Howard, M. J. Bevan, and F. R. Carbone, 1994. T cell receptor antagonist peptides induce positive selection. Cell 76:17–27.

5. Robbins, P. F., Y. F. Li, M. El-Gamil, Y. Zhao, J. A. Wargo, Z. Zheng, H. Xu, R. A. Morgan, S. A. Feldman, L. A. Johnson, A. D. Bennet, S. M. Dunn, T. M. Mahon, B. K. Jakobsen, and S. A. Rosenberg, 2008. Single and dual amino acid substitutions in TCR CDRs can enhance antigen-specific T cell functions. J. Immunol. 180:6116–6131.

6. Depoil, D., and M. L. Dustin, 2014. Force and affinity in ligand discrimination by the TCR. Trends Immunol. 35:597–603.

7. Hivroz, C., and M. Saitakis, 2016. Biophysical aspects of T lymphocyte activation at the immune synapse. Front. Immunol. 7:46.

8. Upadhyaya, A., 2017. Mechanosensing in the immune response. Semin. Cell Dev. Biol. 71:137–145.

9. Cai, E., K. Marchuk, P. Beemiller, C. Beppler, M. G. Rubashkin, V. M. Weaver, A. Gerard, T.-L. Liu, B.-C. Chen, E. Betzig, F. Bartumeus, and M. F. Krummel, 2017. Visualizing dynamic microvillar search and stabilization during ligand detection by T cells. Science 356:eaal3118.

10. Liu, B., W. Chen, B. D. Evavold, and C. Zhu, 2014. Accumulation of dynamic catch bonds between TCR and agonist peptide-MHC triggers T cell signaling. Cell 157:357–368.

11. Das, D. K., Y. Feng, R. J. Mallis, X. Li, D. B. Keskin, R. E. Hussey, S. K. Brady, J. H. Wang, G. Wagner, E. L. Reinherz, and M. J. Lang, 2015. Force-dependent transition in the T-cell receptor β-subunit allosterically regulates peptide discrimination and pMHC bond lifetime. Proc. Natl. Acad. Sci. U.S.A. 112:1517–1522.

12. Sage, P. T., L. M. Varghese, R. Martinelli, T. E. Sciuto, M. Kamei, A. M. Dvorak, T. A. Springer, A. H. Sharpe, and C. V. Carman, 2012. Antigen recognition is facilitated by invadosome-like protrusions formed by memory/effector T cells. J. Immunol. 1102594.

13. Jung, Y., I. Riven, S. W. Feigelson, E. Kartvelishvily, K. Tohya, M. Miyasaka, R. Alon, and G. Haran, 2016. Three-dimensional localization of T-cell receptors in relation to microvilli using a combination of superresolution microscopies. Proc. Natl. Acad. Sci. U.S.A. 113:E5916–E5924.

14. Kim, H.-R., Y. Mun, K.-S. Lee, Y.-J. Park, J.-S. Park, J.-H. Park, B.-N. Jeon, C.-H. Kim, Y. Jun, Y.-M. Hyun, M. Kim, S.-M. Lee, C.-S. Park, S.-H. Im, and C.-D. Jun, 2018. T cell microvilli constitute immunological synaptosomes that carry messages to antigen-presenting cells. Nat. Commun. 9:3630.

15. Razvag, Y., Y. Neve-Oz, J. Sajman, M. Reches, and E. Sherman, 2018. Nanoscale kinetic segregation of TCR and CD45 in engaged microvilli facilitates early T cell activation. Nat. Commun. 9:732.

16. Pullen III, R. H., and S. M. Abel, 2017. Catch bonds at T cell interfaces: impact of surface reorganization and membrane fluctuations. Biophys. J. 113:120–131.

17. Huang, J., V. I. Zarnitsyna, B. Liu, L. J. Edwards, N. Jiang, B. D. Evavold, and C. Zhu, 2010. The kinetics of two-dimensional TCR and pMHC interactions determine T-cell responsiveness. Nature 464:932–936.

18. Hong, J., S. P. Persaud, S. Horvath, P. M. Allen, B. D. Evavold, and C. Zhu, 2015. Force-regulated *in situ* TCR-peptide-bound MHC class II kinetics determine functions of CD4+ T cells. J. Immunol. 195:3557–3564.

19. Feng, Y., K. N. Brazin, E. Kobayashi, R. J. Mallis, E. L. Reinherz, and M. J. Lang, 2017. Mechanosensing drives acuity of αβ T-cell recognition. Proc. Natl. Acad. Sci. U.S.A. 114:E8204–E8213.

20. Sibener, L. V., R. A. Fernandes, E. M. Kolawole, C. B. Carbone, F. Liu, D. McAffee, M. E. Birnbaum, X. Yang, L. F. Su, W. Yu, S. Dong, M. H. Gee, K. M. Jude, M. M. Davis, J. T. Groves, W. A. Goddard III, J. R. Heath, B. D. Evavold, R. D. Vale, and K. C. Garcia, 2018. Isolation of a structural mechanism for uncoupling T cell receptor signaling from peptide-MHC binding. Cell 174:672–687.

21. Hong, J., C. Ge, P. Jothikumar, B. Liu, K. Bai, K. Li, W. Rittase, M. Shinzawa, Y. Zhang, A. Palin, P. Love, X. Yu, K. Salaita, B. D. Evavold, A. Singer, and C. Zhu, 2018. A TCR mechanotransduction signaling loop induces negative selection in the thymus. Nat. Immunol. 19:1379.

22. Feng, Y., E. L. Reinherz, and M. J. Lang, 2018. *αβ* T Cell Receptor Mechanosensing Forces out Serial Engagement. Trends Immunol. 39:569–609.

23. Majstoravich, S., J. Zhang, S. Nicholson-Dykstra, S. Linder, W. Friedrich, K. A. Siminovitch, and H. N. Higgs, 2004. Lymphocyte microvilli are dynamic, actin-dependent structures that do not require Wiskott-Aldrich syndrome protein (WASp) for their morphology. Blood 104:1396–1403.

24. Fisher, P. J., P. A. Bulur, S. Vuk-Pavlovic, F. G. Prendergast, and A. B. Dietz, 2008. Dendritic cell microvilli: a novel membrane structure associated with the multifocal synapse and T-cell clustering. Blood 112:5037–5045.

25. Birnbaum, M. E., R. Berry, Y.-S. Hsiao, Z. Chen, M. A. Shingu-Vazquez, X. Yu, D. Waghray, S. Fischer, J. McCluskey, J. Rossjohn, W. Thomas, and K. C. Garcia, 2014. Molecular architecture of the αβ T cell receptor-CD3 complex. Proc. Natl. Acad. Sci. U.S.A. 111:17576–17581.

26. Grakoui, A., S. K. Bromley, C. Sumen, M. M. Davis, A. S. Shaw, P. M. Allen, and M. L. Dustin, 1999. The immunological synapse: A molecular machine controlling T cell activation. Science 285:221–227.

27. Casal, A., C. Sumen, T. E. Reddy, M. S. Alber, and P. P. Lee, 2005. Agent-based modeling of the context dependency in T cell recognition. J. Theor. Biol. 236:376–391.

28. O’Donoghue, G. P., R. M. Pielak, A. A. Smoligovets, J. J. Lin, and J. T. Groves, 2013. Direct single molecule measurement of TCR triggering by agonist pMHC in living primary T cells. eLife 2:e00778.

29. Wofsy, C., D. Coombs, and B. Goldstein, 2001. Calculations show substantial serial engagement of T cell receptors. Biophys. J. 80:606–612.

30. Favier, B., N. J. Burroughs, L. Wedderburn, and S. Valitutti, 2001. TCR dynamics on the surface of living T cells. Int. Immunol. 13:1525–1532.

31. Springer, T. A., 1990. Adhesion receptors of the immune system. Nature 346:425.

32. Bell, G. I., 1978. Models for the specific adhesion of cells to cells. Science 200:618–627.

33. Pereverzev, Y. V., O. V. Prezhdo, M. Forero, E. V. Sokurenko, and W. E. Thomas, 2005. The two-pathway model for the catch-slip transition in biological adhesion. Biophys. J. 89:1446–1454.

34. Burkhardt, J. K., E. Carrizosa, and M. H. Shaffer, 2008. The actin cytoskeleton in T cell activation. Annu. Rev. Immunol. 26:233–259.

35. Liu, Y., L. Blanchfield, V. P.-Y. Ma, R. Andargachew, K. Galior, Z. Liu, B. Evavold, and K. Salaita, 2016. DNA-based nanoparticle tension sensors reveal that T-cell receptors transmit defined pN forces to their antigens for enhanced fidelity. Proc. Natl. Acad. Sci. U.S.A. 113:5610–5615.

36. Siller-Farfán, J. A., and O. Dushek, 2018. Molecular mechanisms of T cell sensitivity to antigen. Immunol. Rev. 285:194–205.

37. Courtney, A. H., W.-L. Lo, and A. Weiss, 2018. TCR Signaling: Mechanisms of Initiation and Propagation. Trends Biochem. Sci. 43:108–123.

38. Pettmann, J., A. M. Santos, O. Dushek, and S. J. Davis, 2018. Membrane ultrastructure and T cell activation. Front. Immunol. 9:2152.

