## Supporting material for "Mechanical feedback enables catch bonds to selectively stabilize scanning microvilli at T-cell surfaces"

Robert H. Pullen III and Steven M. Abel

### 1 Methods: Computational details

The dynamics of the system are described by a discrete-time, continuous-space stochastic algorithm. This choice was physically motivated by describing the microvillus position as a function of time. The time step ($\Delta t$) was chosen to be sufficiently small so that the discrete-time algorithm provides a good approximation of the underlying continuous-time process. During each time step, the algorithm allows for diffusive hops of particles, binding of TCRs and pMHCs, and dissociation of bonds. At the end of each time step, the position of the microvillus is updated in accordance with its velocity, which impacts the state of the system by changing the lengths of TCR-pMHC bonds and the positions of TCRs relative to the antigen-presenting surface.

Unbound pMHC molecules and TCRs diffuse on the antigen-presenting surface and microvillus tip, respectively. In time interval $\Delta t$, the probability that a given particle attempts a diffusive move is

$$P_{\text{diff}} = 1 - e^{-4D\Delta t/\Delta r^2} , \tag{1}$$

where $D$ is the diffusion coefficient and $\Delta r$ is the displacement. The diffusive step consists of the particle moving a distance $\Delta r$ at a randomly generated angle. The move is accepted if the new position does not overlap with other particles and remains within the appropriate boundaries; otherwise it is rejected and the particle stays at the original location.

The binding of a TCR with a pMHC is governed by an intrinsic on-rate that decays like a Gaussian as the distance varies from the natural length of the TCR-pMHC complex. The total probability of binding for a particular TCR is

$$P_{\text{bind}} = 1 - \exp\left(-\Delta t \sum_{i=1}^{n_{\text{pMHC}}} k^{\text{on}} e^{-(L_i - z_{\text{bond}})^2/2\sigma^2}\right) , \tag{2}$$

where $n_{\text{pMHC}}$ is the number of unbound pMHC molecules within binding distance, $k^{\text{on}}$ is the 2D on-rate for the TCR-pMHC reaction, and $L_i$ is the distance between the TCR and the $i^{\text{th}}$ pMHC molecule. If a binding reaction occurs, the pMHC is chosen with a probability proportional to its individual on-rate with the TCR. We set $\sigma = 5$ nm and restrict the maximum distance for binding between two particles to be twice the natural length of the TCR-pMHC complex ($2z_{\text{bond}}$). The probability that a TCR-pMHC complex dissociates within the time interval $\Delta t$ is

$$P_{\text{diss}} = 1 - e^{-k_{\text{off}}(f)\Delta t} , \tag{3}$$

where $k_{\rm off}(f)$ is the ligand-dependent off-rate.

Upon completion of all diffusive and binding processes, the TCR positions and microvillus tip are updated by moving the microvillus a distance of $V_{\rm MV}(t)\Delta t$ in the $x$-direction. The forces, microvillus velocity, and rates for binding and dissociation reactions are then evaluated given the new configuration. This process is repeated until the final simulation time point is reached.

### 2 Methods: Parameterization of dissociation rates

| | $k_0$ ($s^{-1}$) | $f_0$ (pN) | $k_c$ ($s^{-1}$) | $f_c$ (pN) | $k_s$ ($s^{-1}$) | $f_s$ (pN) |
|---|---|---|---|---|---|---|
| OVA | — | — | 4.241 | 3.150 | 0.374 | 9.280 |
| VSV8 | — | — | 40.00 | 2.286 | 0.050 | 9.413 |
| slip | 2.514 | 5.533 | — | — | — | — |
| strong slip | 0.303 | 5.533 | — | — | — | — |

Table 1: Parameterization of TCR lifetime data

### 3 Functional form of $V_{\rm MV}$

Results obtained with different forms of $V_{\rm MV}$ are shown in Figure S4. We considered two variations of the velocity profile used in the paper. In the first, we increased and decreased the value of the force threshold $f_{\rm MV}$ by a factor of two. Increasing the threshold decreases the height of the peak near 0 $\mu$m/min in the 100% VSV8 system, while narrowing and shifting the range of observed velocities toward $V_0$ in the 100% slip pMHC case. Similarly, decreasing the threshold increases the height of the peak near 0 $\mu$m/min in the 100% VSV8 pMHC system, while broadening and shifting the range of observed velocities toward a lower velocity in the 100% slip pMHC case. Because the peak lifetime of OVA occurs at a lower value of the force and for VSV8, decreasing $f_{MV}$ would lead to systems with OVA behaving more like VSV8 in the paper.

We also considered velocity profiles with a Hill-like (sigmoidal) form:

$$V_{\rm MV} = V_0 \left( \frac{1}{1 + \left( -f_x/F_{\rm MV} \right)^{n_{\rm H}}} \right) \tag{4}$$

for $f_x < 0$ and $V_{\rm MV} = V_0$ for $f_x > 0$. Here, $F_{\rm MV}$ is the Hill reference force, $n_{\rm H}$ is the Hill exponent, and the microvillus velocity is constrained between 0 and $V_0$. Figure S4 shows results from simulations of systems containing nonstimulatory pMHC, OVA pMHC, and VSV8 pMHC. We considered values of $F_{\rm MV} = 25$ pN and $n_{\rm H} = 4$. For both OVA and VSV8, the distributions of velocities are similar given a linear or Hill-like velocity function. However, the distribution of velocities for nonstimulatory pMHC has a significant peak near $V_0$ with the Hill function. This is consistent with the net horizontal forces on the microvillus tip being typically less than 25 pN. Thus, the Hill function results in an even more pronounced impact of catch bonds due to the inability of nonstimulatory ligands to significantly slow the microvillus tip on their own.

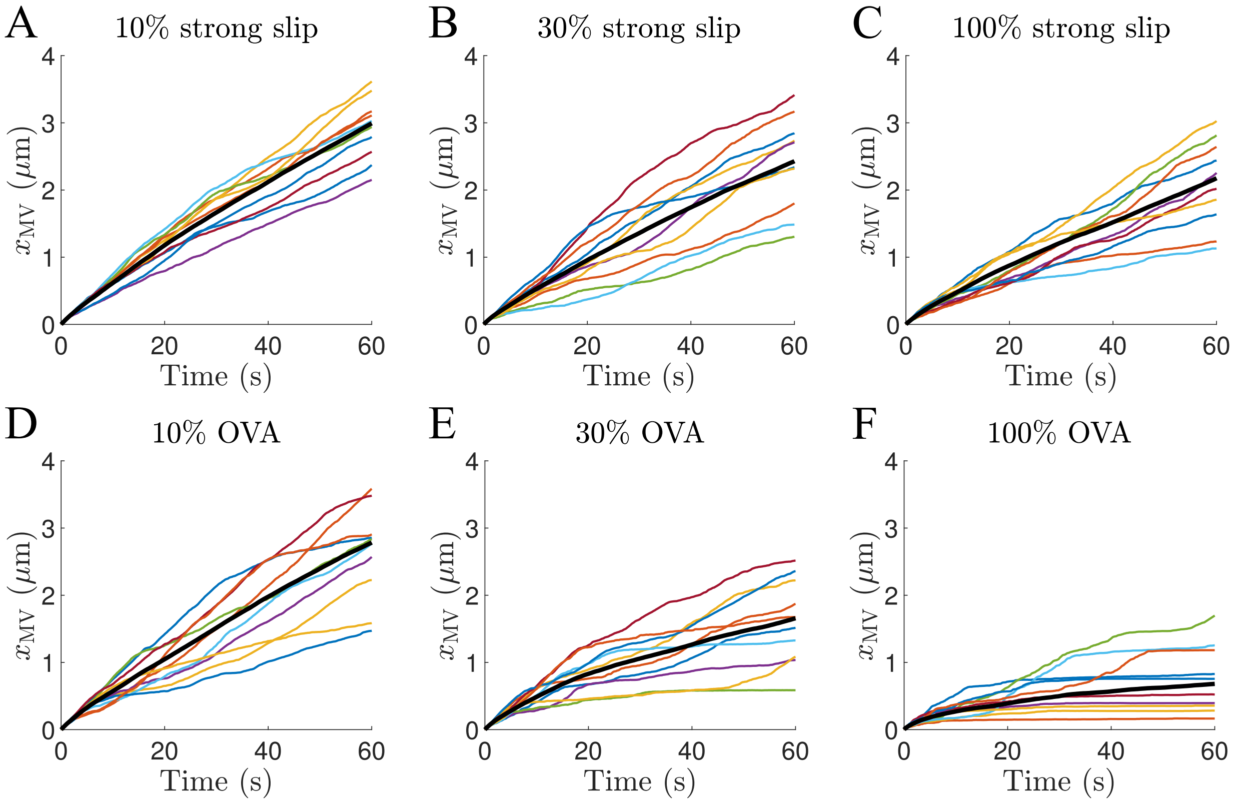


Figure 1: Displacements of microvilli for different fractions of strong-slip (A - C) and OVA (D - F) pMHC. Black lines show the average microvillus displacement calculated from 25 independent trajectories. Colored lines show the displacement of individual microvilli (10 shown).

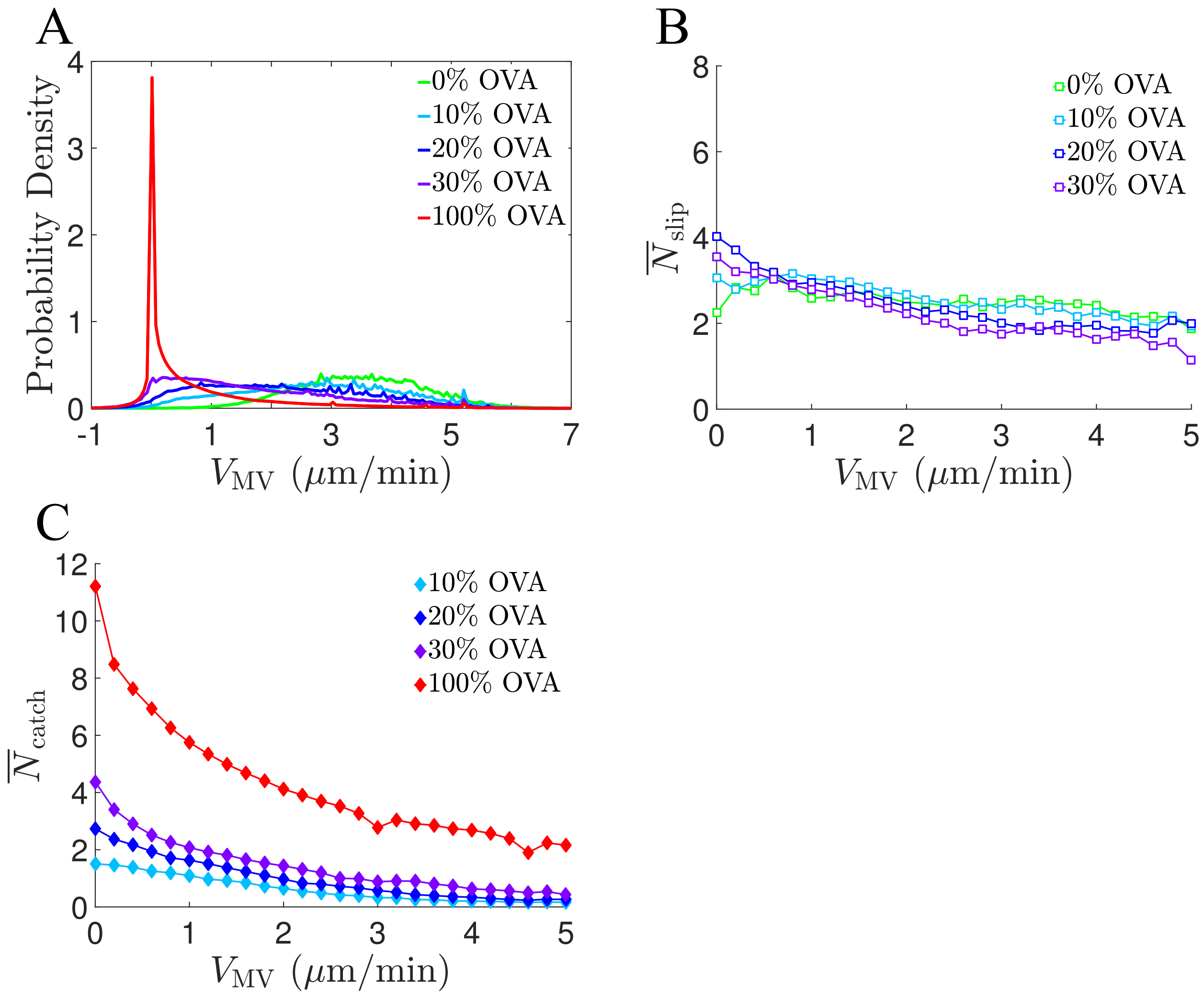


Figure 2: (A) Probability density of the microvillus velocity at various fractions of OVA pMHC. (B, C) The average number of slip and catch bonds as a function of the microvillus velocity for the OVA system.

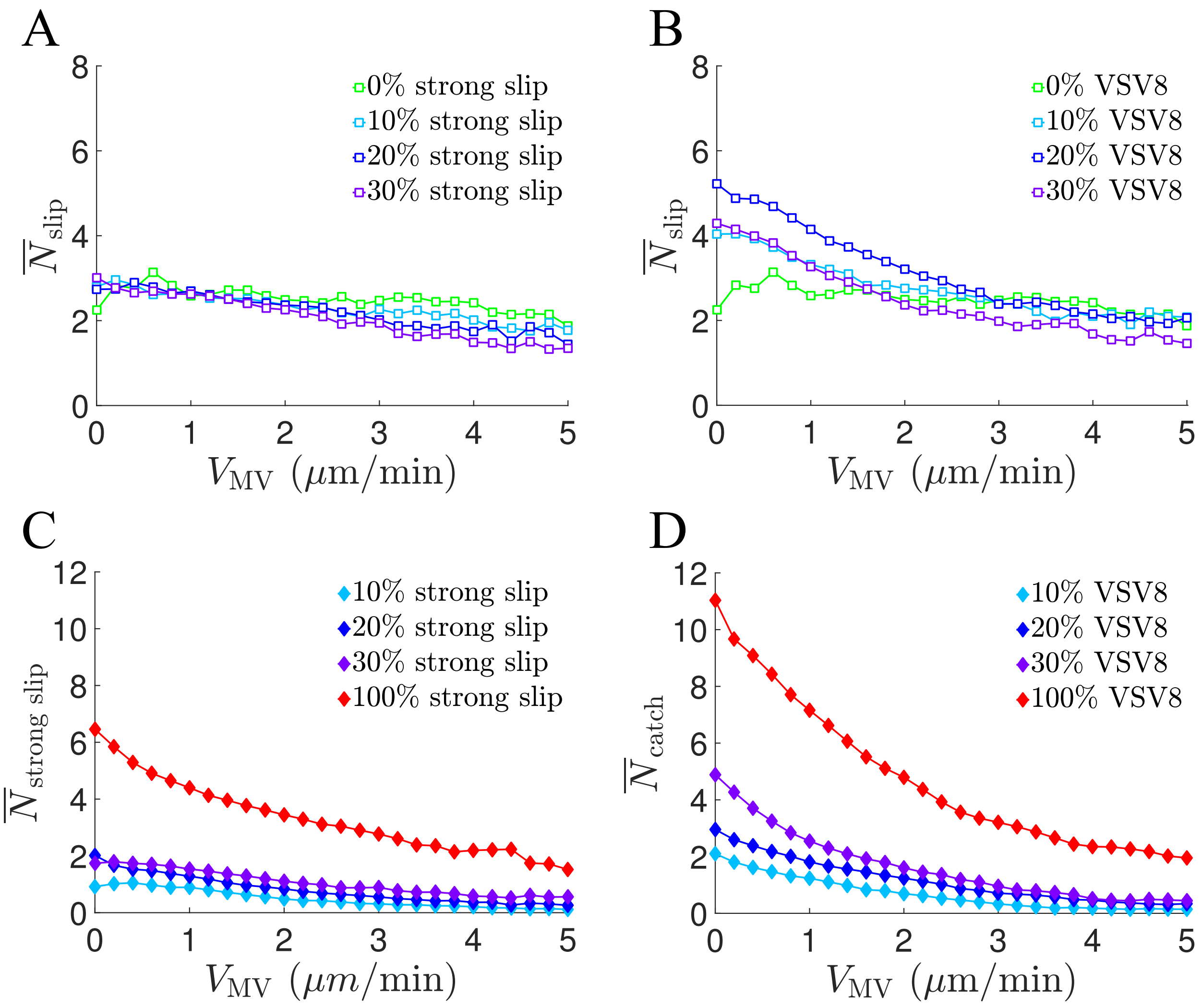


Figure 3: (A, B) The average number of nonstimulatory slip bonds as a function of the microvillus velocity for varying fractions of strong-slip and VSV8 pMHC. (C, D) The average number of strong-slip and catch bonds for the same systems.

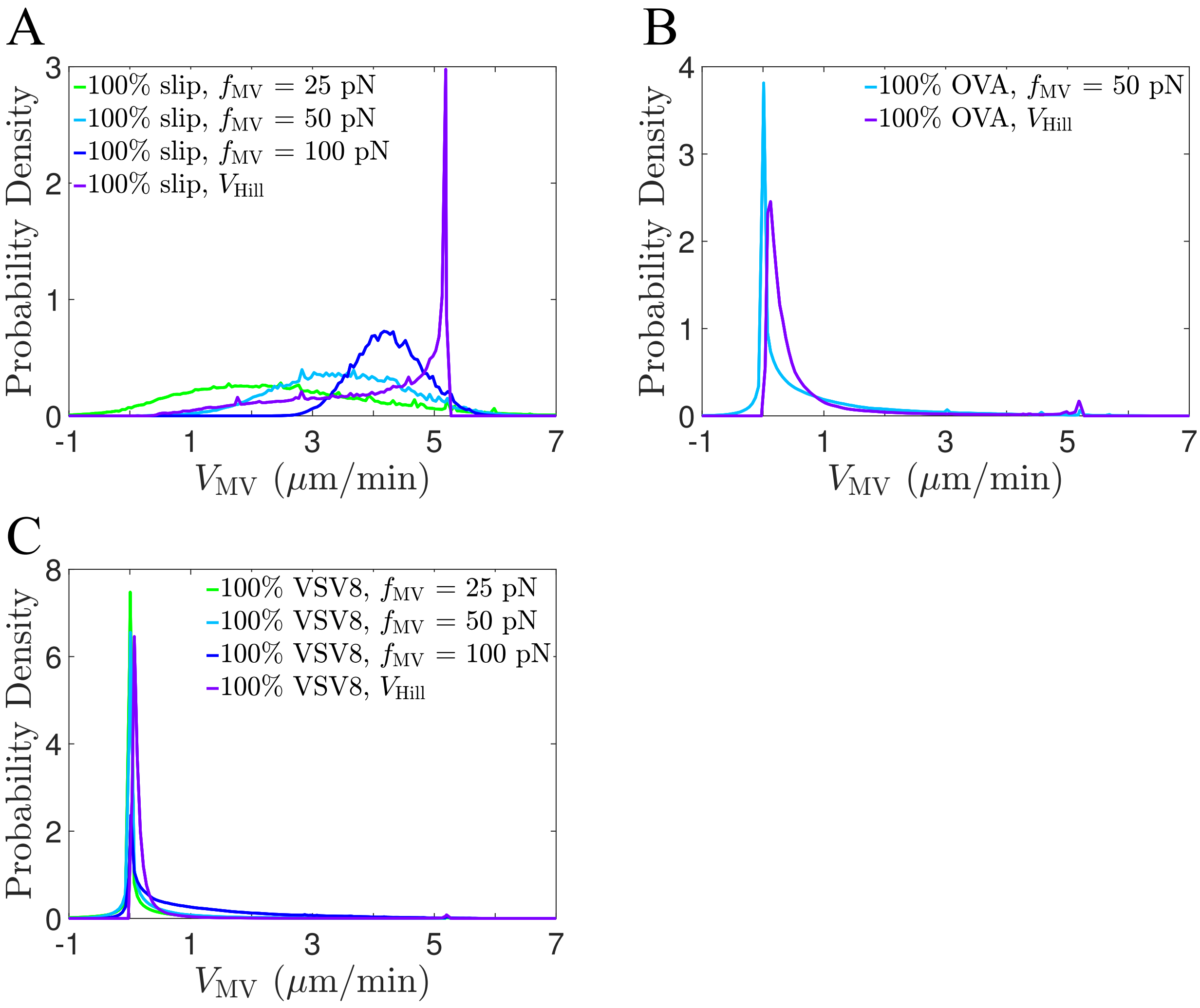


Figure 4: Characterization of the velocity distribution for different forms of $V_{\rm MV}$ with 100% slip (A), 100% OVA (B), and 100% VSV8 (C). The linear response is considered with $f_{\rm MV} = 25$ pN and 100 pN (compared with $f_{\rm MV} = 50$ pN in the main text). The Hill-like response is considered with $F_{\rm MV} = 25$ pN and $n_{\rm H} = 4$.

| **A** | | 0% VSV8 | 10% VSV8 | 20% VSV8 | 30% VSV8 | 100% VSV8 |
|---|---|---|---|---|---|---|
| $V_{\mathrm{MV}}$ ($\mu$m/min) | | $3.486 \pm 1.030$ | $2.232 \pm 0.781$ | $1.346 \pm 0.555$ | $0.948 \pm 0.426$ | $0.230 \pm 0.083$ |
| $\tau$ (s) | slip | $0.117 \pm 0.110$ | $0.141 \pm 0.157$ | $0.167 \pm 0.193$ | $0.178 \pm 0.213$ | — |
| | catch | — | $0.712 \pm 1.022$ | $0.671 \pm 1.153$ | $0.549 \pm 1.261$ | $0.417 \pm 1.045$ |
| $N_{\mathrm{bonds}}$ | slip | $2.361 \pm 1.452$ | $2.866 \pm 1.811$ | $4.037 \pm 2.540$ | $3.568 \pm 2.116$ | — |
| | catch | — | $0.804 \pm 0.901$ | $1.837 \pm 1.446$ | $3.316 \pm 2.467$ | $10.255 \pm 3.890$ |

| **B** | | 0% OVA | 10% OVA | 20% OVA | 30% OVA | 100% OVA |
|---|---|---|---|---|---|---|
| $V_{\mathrm{MV}}$ ($\mu$m/min) | | (see above) | $2.785 \pm 0.862$ | $2.148 \pm 0.668$ | $1.658 \pm 0.521$ | $0.681 \pm 0.313$ |
| $\tau$ (s) | slip | — | $0.127 \pm 0.128$ | $0.140 \pm 0.149$ | $0.153 \pm 0.170$ | — |
| | catch | — | $0.445 \pm 0.355$ | $0.475 \pm 0.402$ | $0.530 \pm 0.500$ | $0.603 \pm 0.614$ |
| $N_{\mathrm{bonds}}$ | slip | — | $2.448 \pm 1.540$ | $2.498 \pm 1.652$ | $2.542 \pm 1.719$ | — |
| | catch | — | $0.514 \pm 0.693$ | $1.098 \pm 1.114$ | $1.978 \pm 1.837$ | $8.541 \pm 4.507$ |

| **C** | | 0% strong slip | 10% strong slip | 20% strong slip | 30% strong slip | 100% strong slip |
|---|---|---|---|---|---|---|
| $V_{\mathrm{MV}}$ ($\mu$m/min) | | (see above) | $2.993 \pm 0.880$ | $2.738 \pm 0.852$ | $2.430 \pm 0.787$ | $2.176 \pm 0.651$ |
| $\tau$ (s) | slip | — | $0.122 \pm 0.118$ | $0.127 \pm 0.128$ | $0.133 \pm 0.138$ | — |
| | strong slip | — | $0.384 \pm 0.263$ | $0.414 \pm 0.310$ | $0.439 \pm 0.342$ | $0.466 \pm 0.366$ |
| $N_{\mathrm{bonds}}$ | slip | — | $2.179 \pm 1.372$ | $2.128 \pm 1.378$ | $2.077 \pm 1.392$ | — |
| | strong slip | — | $0.364 \pm 0.601$ | $0.694 \pm 0.845$ | $1.052 \pm 0.982$ | $3.479 \pm 2.011$ |

Table 2: Average $\pm$ SD for the microvillus velocity, bond lifetimes, and number of bonds for systems with VSV8 (A), OVA (B), and strong slip (C). All values are calculated from 25 independent trajectories.
